# Validation of a combined cylindrical shield and partial-coverage mobile OPM system for detecting neuromagnetic sensorimotor responses in humans

**DOI:** 10.1101/2025.06.10.657831

**Authors:** Lyam M. Bailey, Timothy Bardouille

**Affiliations:** Department of Physics & Atmospheric Science, Dalhousie University, Halifax, NS B3H4R2, Canada

## Abstract

Optically pumped magnetometers (OPMs) have emerged as a promising technology for neuromagnetic recording in humans. Current state-of-the-art OPM systems are housed in immobile magnetically-shielded rooms to reduce external electromagnetic noise, and typically comprise sensor arrays covering the entire head. Here we sought to validate a low-cost, mobile OPM system comprising a small cylindrical mu-metal shield and partial sensor coverage. Twelve participants underwent right-sided median nerve stimulation (MNS) and cued right-handed button-pressing intended to elicit ubiquitous sensorimotor responses: somatosensory-evoked fields (SEFs; comprising N20m, P35m and P60m components) and event-related (de)synchronisation (ERD/ERS) of oscillatory neuronal rhythms in the mu and beta frequency ranges. Following MNS, we observed robust N20m and P60m peaks, as well as the expected mu ERD and beta ERS effects. Moreover, we successfully localized these responses to expected cortical generators using distributed source modelling. SEFs and mu ERD were both maximal in left (i.e., contralateral to stimulation) primary somatosensory cortex (central sulcus, postcentral gyrus and sulcus), while beta ERS appeared more anteriorly, in the central sulcus and precentral gyrus. By contrast, results from the button-pressing paradigm were less conclusive—we observed beta ERS (but not mu/beta ERD), and an atypical distribution of the ERS effect over posterior ROIs. Overall, our findings provide proof-of-principle support for the use of our system in the context of passive (e.g., MNS) paradigms; its viability for cued movement tasks will require further development. Based on these results, we make recommendations for further developments in mobile and partial-coverage OPM.

---

Over the past decade, optically-pumped magnetometers (OPMs) have emerged as a promising alternative to conventional systems used for magnetoencephalography (MEG)—a non-invasive neuroimaging technique for neuroelectrophysiological recording in humans. MEG is a valuable tool for neuroscientists because it allows us to record millisecond-to-millisecond fluctuations in neuronal activity (e.g., over the course of rapidly changing experimental conditions), and offers relatively high (2-5 mm) spatial resolution on the cortical surface (Brookes et al., 2022; Fred et al., 2022). Conventional cryogenic MEG systems, which rely on superconducting quantum interference devices (SQUIDs), are relatively inaccessible for most research groups because they are expensive, require housing in bespoke facilities, and demand a constant supply of liquid helium to keep their sensors operational. Moreover, SQUID-MEG is extremely sensitive to participant movement; this limits its utility in individuals who have difficulty remaining still throughout a scan (e.g., children and some clinical populations). By contrast, OPM-MEG systems are less costly to purchase, do not require liquid helium, and are relatively insensitive to movement (see Brookes et al., 2022 for review and discussion), while also providing comparable or superior sensitivity to human neuronal activity (Borna et al., 2020; Boto et al., 2017; Marhl et al., 2022; Rier et al., 2024).

Despite the gains (both financial and practical) afforded by OPM-MEG, some barriers to accessibility remain. In particular, these systems are often placed inside magnetically shielded rooms (MSRs) to mitigate external sources of noise (e.g., see Holmes et al., 2022). These MSRs inherently limit the mobility of OPM-MEG and take up large amounts of physical space; this in turn limits its translation to clinical and rural environments. Moreover, while OPM-MEG is relatively inexpensive compared to SQUID-MEG, procuring enough OPM sensors for whole-head coverage (typical in MEG) requires substantial financial investment—often hundreds of thousands of dollars. Efforts to overcome these barriers will help improve the utility and accessibility of OPM as a tool for experimental and clinical neuroscience.

Here we evaluate a low-cost mobile OPM system, previously described in Bardouille et al. (2024), for detecting neuromagnetic responses in humans. In brief, this system comprises a dual-wall cylindrical mu-metal shield, mounted on a moveable wooden cradle, and a rigid helmet in which OPM sensors can be positioned against a participant’s head (Figure 1a). The system also comprises sixteen single-axis OPM sensors which can be flexibly arranged to overlay a given area of the head. In this work for example, we arranged sensors over left sensorimotor areas, with a view to capture neuromagnetic signals elicited by contralateral (right-sided) stimulation and movement (Figure 1b).

**Figure 1.**
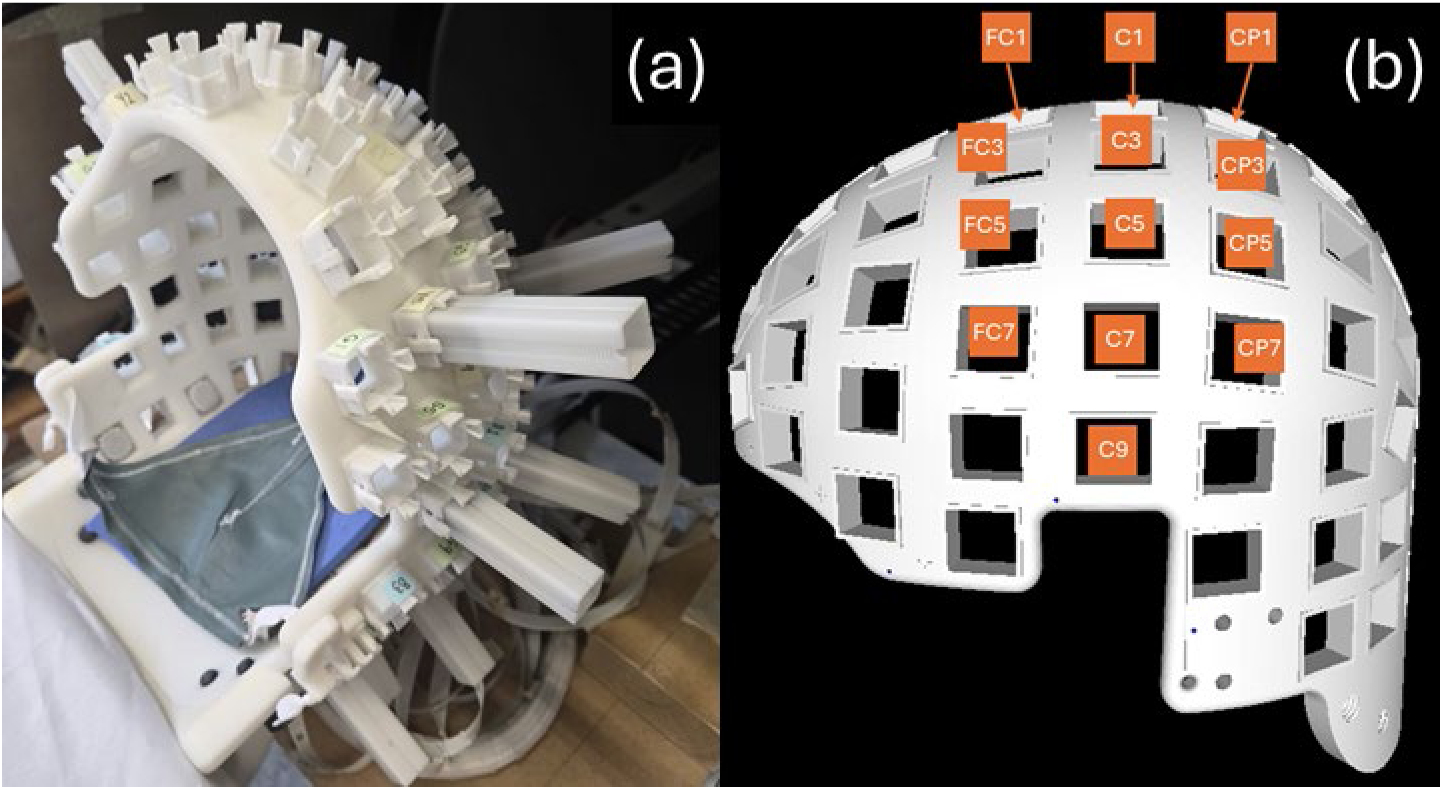
The mobile OPM system. (**a**) The rigid helmet into which sensors may be inserted. (**b**) The sensor configuration used in this study.

Voluntary movement and median nerve stimulation (MNS) reliably elicit neuromagnetic biomarkers of sensorimotor processing. In particular, MNS elicits a somatosensory evoked field (SEF) complex that is characterised by peaks around 20, 35, and 60 ms following stimulation (labelled N20m, P35m, and P60m respectively). Evidence from MEG indicates that the SEF originates in primary somatosensory cortex, in Brodmann area 3b (i.e., the posterior bank of the central sulcus) contralateral to the side of stimulation (Kimura & Hashimoto, 2001; Lin et al., 2005; Ou et al., 2009; Wikström et al., 1996). In addition, MNS and voluntary movement both elicit neuronal rhythm changes in the mu (8-15 Hz) and beta (15-30 Hz) frequency ranges. These changes entail an immediate reduction in mu and beta power, relative to a preceding baseline period (event-related desynchronization; ERD); this is followed by a return to baseline levels in the mu range and a “rebound” (event-related synchronisation; ERS) in beta power (Bardouille et al., 2019; Byrne et al., 2017; Houdayer et al., 2006; Jurkiewicz et al., 2006; Pfurtscheller et al., 1996). In terms of their underlying cortical generators, mu and beta ERD are typically localized to central-to-posterior sensorimotor cortex contralateral to movement or stimulation, while the beta rebound tends to appear more anteriorly (Bardouille et al., 2019; Jurkiewicz et al., 2006). SEFs and oscillatory rhythm changes are ubiquitous in MEG research and can be detected by contemporary OPM systems housed in MSRs (An et al., 2022, 2024; Hill et al., 2020; Rier et al., 2024; Schofield et al., 2024). As such, they provide a useful benchmark by which to validate our system.

We evaluated the ability of our OPM-MEG system to detect these ubiquitous sensorimotor responses following MNS and voluntary movement. Besides detecting these responses at the sensor level, an important quality assurance metric for any MEG system is source localization accuracy. This issue is particularly salient for evaluating our system, considering that low sensor count and partial head coverage might impede localization accuracy. As such, we assessed whether SEF and oscillatory responses detected at the sensor level could be localized to areas of contralateral sensorimotor cortex consistent with the wider literature. Previous work with this (Bardouille et al., 2024) and similar (Borna et al., 2017, 2020; Boto et al., 2017) systems with low channel counts has relied on simple dipole fitting for source localization. As a secondary aim, we investigated the applicability of distributed source modelling, which (to our knowledge) is typically applied to data from high-coverage (∼90+ channels) systems. Unlike dipole fitting, distributed source modelling is informative as to the spatial extent of cortical responses; establishing its viability here would be valuable for future research using sparse, partial-coverage systems such as this one.

Participants completed two right-sided MNS protocols, as well as a right-handed cued button pressing task, inside the mobile scanner. We expected that MNS and movement would elicit typical modulations in the mu and beta ranges (mu and beta ERD followed by beta ERS), while MNS should also induce SEFs. Moreover, we expected that distributed source modelling would localize these responses to contralateral sensorimotor cortex. More precisely: SEFs and mu/beta ERD should be maximal in the central sulcus and postcentral gyrus. Meanwhile, beta ERS should be maximal more anteriorly, in the precentral gyrus. Confirmation of these predictions would represent an important validation step for our low-cost, mobile system as a viable alternative to MSR-based MEG.

## Methods

### Subjects

Twelve volunteers (6 female, aged 19-36, mean age 24.17) were recruited through on-campus advertising at Dalhousie University and were compensated with $25 CAD for their time. All participants were right-handed^1^ as determined by the Edinburgh Handedness Inventory (Oldfield, 1971), neurologically healthy (no previous stroke or brain lesion), were not taking medication for a neurological or psychiatric disorder, and were free of metal implants, hearings aids, and non-removable metal piercings. Three subjects had non-removable wire dental retainers; these were de-magnetised immediately prior to OPM scanning using an audio/video tape eraser (RadioShack, Canada). All procedures were approved by the Dalhousie University Research Ethics Board (REB #2021-5586). One participant’s data from the cued button pressing task was discarded due to excessive motion; we therefore report data from 12 and 11 participants, from the MNS and button pressing tasks respectively.

### Stimuli and Apparatus

The following two sections (“Cylindrical mu-metal shield” and “OPM sensor array”) have been adapted from Bardouille et al. (2024).

#### Cylindrical mu-metal shield

The cylindrical shield comprises inner and outer layers of mu-metal, with 43 cm and 48 cm radii and axial lengths of 1.9 m and 2.1 m, respectively (American Magnetics, Oak Ridge, TN, USA). The open end of the cylinder (for participant insertion) includes a 45 cm long reducer to a radius of 38 cm. The opposite ends of the inner and outer shields are closed with friction-fitted end caps. The thickness of the 3 mu-metal components is 1.8 mm. A smaller-diameter polymer cylinder includes coils for nulling the homogenous field along each cardinal axis (i.e., active shielding; not used in this study). The cylinder is mounted on a wooden cradle, and participants are supported (lying supine) on a retractable wooden bed, on a non-metal rail system which allows the bed to be inserted into the cylinder.

#### OPM sensor array

Sixteen FieldLine v2 single-axis OPM sensors (Fieldline Inc., Boulder, CO, USA), with less than 20 fT/√Hz sensitivity and a bandwidth of 0-150 Hz, were used in this study. Device control and 24-bit data acquisition were managed via vendor-supplied v2 electronics chassis and a personal computer running vendor-supplied acquisition software (Recorder, v1.6.7) on the Windows 11 operating system. A rigid helmet for mounting OPM sensors is fixed to the end of the wooden bed (Figure 1a). OPM sensors are inserted into slots in the helmet at variable depths, such that they can be positioned against a participant’s head. The OPM sensor positions with respect to the helmet can be precisely known by measuring the depth of the sensor in each slot. The orientation is precisely defined for each slot based on the helmet manufacture, independent of depth. Three of the 16 sensors were fixed to the end of the bed (close to the helmet) and arranged in orthogonal orientations to be used as reference sensors; the remaining 13 were inserted into the helmet for participant scans. These 13 sensors were arranged in a left-sided configuration (Figure 1b) over sites where sensorimotor responses ought to be maximal (FC1, C1, CP1, FC3, C3, CP3, FC5, C5, CP5, FC7, C7, CP7, C9).

### Procedures

Each participant completed informed consent and was given a brief tour of the OPM system before beginning the experiment. Prior to the MEG recording, we collected a 3-D digital image of the participant’s head shape using an optical imaging system (EinScan H2, Shining 3D) – to register the MEG data to a template MRI. Coloured markers were placed on the fiducial points (nasion, left and right preauricular points) to facilitate manual identification in the resultant images. Then, the participant was instructed to lie supine on the scanner bed with their head positioned inside the OPM helmet. Thirteen OPM sensors were inserted into the helmet such that they made direct contact with the participant’s head. We collected a second 3-D digital image of the front of the participant’s head positioned inside the OPM helmet – to register the participant’s head to the MEG sensors. Participants completed four scans in a single session, always in the same order: two MNS protocols, a cued button-press task, and a resting-state scan in which they were instructed to lie still with their eyes closed for approximately 5 minutes (data from the resting scan is not reported here). Following the experiment, the depth of each sensor in the helmet was manually recorded.

#### Median nerve stimulation (MNS)

Prior to scanning we fitted each participant with moistened stimulation electrodes housed in a plastic casing, positioned such that the electrodes aligned with the median nerve on their right inner wrist. The electrodes were connected via an insulated cable to a DS7A Constant Current Stimulator (Digitimer, Welwyn Garden City, UK). Stimulation was delivered at 250 V in single 500 µs pulses. Stimulation current was the minimum current required to elicit a small twitch of the right thumb (determined for each participant individually). Participants completed two MNS protocols—labelled “short MNS” and “long MNS”—lasting approximately 5 and 7 minutes respectively. In the first (short MNS) protocol, participants received 300 stimulations delivered 1-1.4 s apart. In the second (long MNS), participants received 80 stimulations delivered 5.6-7.6 s apart. Data from the short and long MNS protocols were used to investigate SEF responses and oscillatory modulations respectively, since longer inter-stimulus intervals (ISIs) are needed to properly quantify beta ERS (Bailey & Bardouille, 2025; Pakenham et al., 2020; Pfurtscheller & Lopes da Silva, 1999).

#### Cued button pressing

Participants heard 80 auditory tones (1000 Hz), delivered 7.0-9.0 s apart, and were instructed to press a button with their right index finger following each tone. Responses were made using an MEG-compatible, non-magnetic button box (Current Designs, Philadelphia, USA) connected by fibre-optic cable to the recording computer. Auditory tones were presented by a non-magnetic speaker (SoundShower, Panphonics, Finland) placed outside of the shield, at a clearly audible sound pressure level. This scan lasted approximately 10 minutes. Data from this scan was used to investigate movement-induced oscillatory changes.

### Data Analysis

Unless otherwise stated, all analyses were performed in the Python environment using functions from the MNE (Gramfort et al., 2013) and Open3D (Zhou et al., 2018) libraries, and custom scripting.

#### Registration of sensor data to a standard head model

We relied on the FreeSurfer template brain for source localization (see Data Analysis); this required co-registration of participants’ head digitization and sensor positions to the template head model. We first conducted surface reconstruction, and computed a boundary element model (BEM; Hamalainen & Sarvas, 1989), from the T1 template image using FreeSurfer’s recon-all algorithm. We then applied the following procedure for each participant.

The two 3D images of the participant’s head, as well as a vendor-supplied computer-aided design image of the OPM helmet, were manually aligned using *MeshLab*. The positions of the fiducial markers were manually determined using the 3D image acquired outside of the helmet. These locations defined the head coordinate frame (x-axis through the preauricular points and y-axis through the nasion). Sensor data were registered to this coordinate frame, as follows. Sensor positions and orientations in the helmet coordinate frame were determined based on known locations of each sensor slot and manually recorded sensor depths. Following this, the 3-D image of the head in the helmet was registered to the head only image using the iterative closest point (ICP) algorithm in Meshlab. We manually identified four points that were common to the head-only and head-in-helmet images (e.g., the three fiducial markers and the tip of the nose); these points were used for ICP. Finally, a vendor-supplied helmet shape was aligned to the sensor slots of the head in helmet 3-D image, using the same ICP procedure. The two resulting transformation matrices were combined with the sensor positions in the helmet coordinate frame to provide the sensor positions in the head coordinate frame. MRI to head registration proceeded as follows. The template anatomical T1 MRI was manually scaled and translated for optimal fit to each individual head, using a visual matching process implemented via the MNE coregistration graphic user interface. Importantly, the template BEM and brain surface were also scaled alongside the T1 image; this allowed us to compute a forward model and source-space grid along the (scaled) cortical surface for each participant.

#### Preprocessing

Raw OPM data was notch filtered to remove line noise and its 1^st^ and 2^nd^ order harmonics (60, 120, and 180 Hz) and then band-pass filtered between 3 and 150 Hz. We removed channels with remaining high-frequency spikes; these were identified by computing a z-scored spectral amplitude per channel in the 120-145 Hz band. We removed channels with z-score > 2.0. We also visually inspected the raw data to identify and remove any additional bad channels (e.g., with repeating artifacts such as extreme amplitude spikes). Signals that were strongly correlated with the three reference sensors were removed using reference array regression as described in Bardouille et al (2024). In brief, we performed a multivariate linear regression which treated each reference sensor as a dependent variable, and each channel as an independent variable. The residuals from each channel were taken as reference-adjusted time courses for that channel and carried through to subsequent analyses. We then performed homogenous field correction (HFC; order=1) to attenuate noise in the raw data arising from ambient magnetic fields with a consistent and homogenous spatial pattern over time (Bardouille et al., 2024; Tierney et al., 2021).

The filtered, re-referenced, and HFC-corrected data was segmented into epochs according to event timing in each scan. Epochs from the short MNS data were 0.7 s in length (-0.2 to +0.5 s relative to stimulation onset); long MNS epochs were 6.4 s (-1.2 to +5.2 s); button press epochs were 7.4 s (-2.2 to +5.2 s relative to button press offset). Prior to epoching, we identified and discarded any erroneous events (i.e., non-cued button presses) from the button-press data, as well as events immediately before and afterwards (un-cued button presses would likely interfere with mu/beta responses from adjacent trials; Bailey & Bardouille, 2025).

Epochs were baseline-corrected by subtracting the mean baseline signal from the entire epoch; baseline periods for each dataset were -0.2 to -0.1 s (short ISI), -1.0 to -0.5 s (long MNS), and -2.0 to -1.0 s (button press). We visually inspected the epoched data to identify and remove artifacts (we rejected 1.39%, 6.35%, and 4.84% of epochs from the short MNS, long MNS, and button-press data respectively). For the short MNS data, we computed the evoked field at each sensor as the inter-trial average. With respect to the long MNS and buttonPress data, we subtracted the inter-trial average from all epochs in each dataset (separately for each participant and channel) to reduce the expression of the evoked response in the time-frequency plot, as analysis of these data focused on ERS and ERD.

#### Source localization

We performed distributed source modelling on the evoked (short MNS) or epoched (long MNS and button-press) data from each task using the same procedure, described as follows. We computed a forward model and source-space grid for each participant, based on their scaled template brain surface and BEM. We then applied minimum norm estimation to project the sensor-level data to the cortical surface. We constrained minimum norm estimation to a relatively large patch of cortex beneath our sensors, comprising a combination of the six regions of interests (ROIs) shown in Figure 2 and described below (applying this constraint greatly reduced processing time and did not affect participant-level evoked responses in these ROIs). This returned a time course for every epoch / evoked field, at every vertex in the constrained source space. Finally, for each epoch / evoked response, we extracted one representative time course as the first principal component of all source estimates within the ROI^2^from each of six a-priori left-hemisphere ROIs. These ROIs were: the superior precentral sulcus, precentral gyrus, central sulcus, postcentral gyrus, postcentral sulcus, and the superior parietal lobule (Figure 2). We selected these ROIs because of their proximity to our sensors, and because they have previously been used to investigate movement-related oscillatory changes (Power & Bardouille, 2021). ROIs were extracted from a standard anatomical parcellation (Destrieux et al., 2010) of the FreeSurfer template brain, morphed to each subject’s cortical surface.

**Figure 2.**
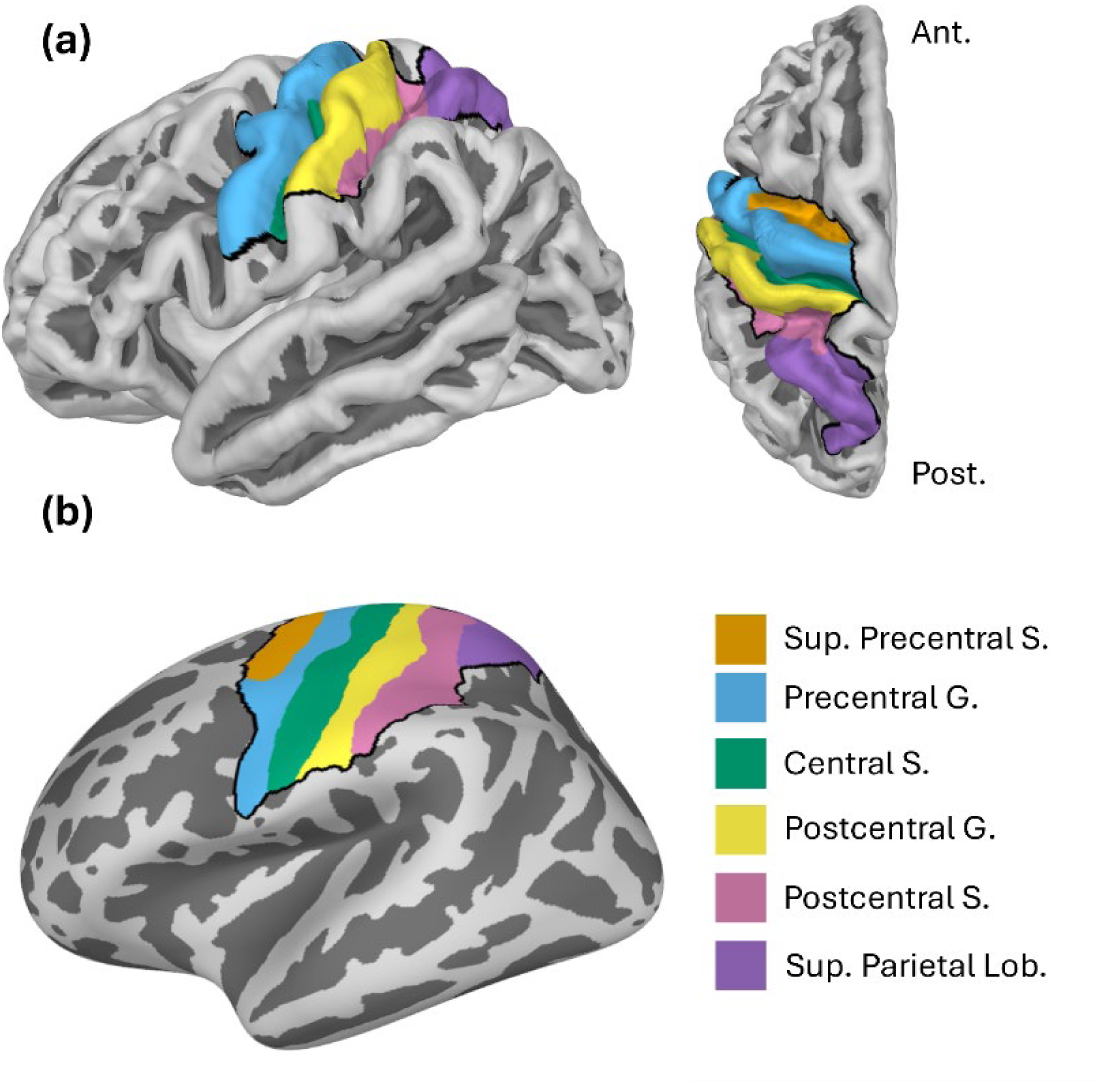
Anatomical regions of interest (ROIs) used for source localization. ROIs are shown on **(a**) a brain surface from lateral (left) and dorsal (right) views, and (**b**) an inflated surface. The dark border shows the area to which source localization was constrained. Abbreviations: *Sup*. Superior; *S*. sulcus; *G*. gyrus.; *Lob*. Lobule.

#### Time-frequency analyses

We conducted time-frequency analyses on the long MNS and button press datasets independently, using the following procedure applied to sensor-level and source-estimated data. We computed time-frequency responses (TFRs) by applying Morlet wavelet analysis to the epoched data (1-40 Hz, 1 Hz steps, number of cycles equal to 1/3 of the center frequency). This produced one TFR array (time x frequency) per epoch and channel/ROI. We removed the first and last 0.2 s from each TFR array to eliminate edge artifacts induced by the wavelet analyses and averaged together the epoch TFRs for each participant and channel/label. Next, we baseline-corrected each TFR array by taking the logarithm of the ratio between power at each time point and mean baseline power, at each frequency (baseline was - 1.0 to -0.5 s for long MNS and -2.0 to -1.0 s for button press). Finally, we averaged the baseline-corrected TFRs over the mu (8-15 Hz) and beta (15-30 Hz) frequency ranges, producing a mu and beta power change timecourse for every participant and channel/ROI. Grand-average TFRs and timecourses were computed by averaging across participants.

#### Statistical analyses

We took a Bayesian approach to verify statistical evidence for SEF peaks and task-induced oscillatory changes relative to baseline amplitude/power. We computed Bayes factors BF_10_ which quantify relative support for the data under H_1_ (which predicts a difference from baseline) versus H_0_ (no difference from baseline). Within this framework, BF_10_ values > 1.0 and < 1.0 signal support for the data under H_1_ and H_0_ respectively; the magnitude of BF_10_ in either direction (i.e., the degree of deviation from 1.0) signals the quantitative strength of evidence (Dienes, 2014, 2016; Lee & Wagenmakers, 2014; Schmalz et al., 2021). Bayes factors offer an alternative to conventional *p* values; they are informative as to quantitative strength of evidence (rather than merely rejecting H_0_), and do not require multiple comparison correction (Dienes, 2016; Teichmann et al., 2022). The following procedures were performed on source-localized timecourses from each of our six ROIs. In the interests of maintaining reasonable scope, we do not report statistical analyses on the sensor-level results.

To quantify SEF peaks in the short MNS data, we defined narrow (3 ms) time windows to capture each peak in participants’ evoked fields for statistical testing. We first defined three a-priori 10 ms windows, centered on 20, 35, and 60 ms following stimulation onset (broadly encompassing the expected N20m, P35m and P60m respectively). Within each window, we then defined a smaller 3 ms window as +/-1 ms around the peak latency in the grand-average (over subjects) evoked field. We also defined a baseline time window as -0.2 to -0.1 s. We then conducted one-tailed^3^ paired Bayesian t-tests to compare mean (over time) amplitude at the N20m, P35m, and P60m time windows to that of the baseline window. Bayesian tests were implemented in the R environment using functions from the BayesFactor package (Morey et al., 2022)^4^. Each t-test returned a Bayes factor BF_10_ wherein values > 1.0 could be interpreted as evidence for a difference in amplitude between the peak and baseline.

To quantify oscillatory changes, we defined a-priori time windows to capture baseline activity, ERD and ERS. For the long MNS data, the baseline period was defined as -1.0 to -0.5 s; the ERD period as 0.2 to 0.4 s; the ERS period as 0.5 to 1.0 s. For the button-press data, these periods were -2.0 to -1.0, -0.25 to 0.25, and 0.5 to 1.0 s respectively. We used a shorter baseline time windows for the long MNS data because this paradigm had a shorter ISI, thus providing less time for the beta rebound to end. Furthermore, the suppression window for the long MNS analysis was delayed to avoid the artifact induced by the MNS current. The window is still centred on the beta ERD peak latency, however, since ERD follows stimulation but slightly precedes movement (see Figure 2, for example). For each participant and dataset, we averaged the computed mu and beta timecourses (see time-frequency analyses) over the baseline, ERD, and ERS time windows. This returned, per participant, a single value representing mean mu or beta power in each time window (with the exception that we did not compute mu power in the ERS periods, since there is no evidence for mu ERS in these contexts). To establish statistical support for mu/beta ERD, we conducted left-sided paired Bayesian t-tests to assess whether mean power during the ERD period was lower than mean power during the baseline period. For beta ERS, we conducted a right-sided t-test to assess whether mean power in the ERS period was higher than that of the baseline period.

#### Quantifying signal-to-noise ratio (SNR)

We computed SNR around the N20m SEF peak at both the sensor and source levels. This enabled straightforward comparison of data quality in our MEG system to that of others, as previous estimates of SNR have been based on the N20m (Antonakakis et al., 2020; Borna et al., 2020; Boto et al., 2017; Kimura & Hashimoto, 2001). Moreover, the N20m was the most statistically robust response observed in our study (see Results); this measure is therefore reflective of maximal data quality attainable from our system.

We computed SNR independently for each subject, sensor, and ROI as the absolute difference in mean amplitude between the baseline window and the (3 ms peak) N20m window, divided by the standard deviation of the baseline window. We then quantified sensor- and source-space SNR for each subject as the maximal value across sensors and ROIs respectively, similar to Antonakakis et al. (2020).

## Results

We investigated somatosensory-evoked fields (SEFs) elicited by an MNS paradigm with relatively short inter-stimulus intervals (labelled “short MNS”), and mu and beta modulation in data from two tasks: an MNS paradigm with relatively long inter-stimulus intervals (labelled “long MNS”) and a cued button-pressing paradigm. We investigated these responses at both the sensor and source levels. Sensor-level results are shown in Figure 3, while source-level results for the short MNS, long MNS, and button press data are shown in Figures 4, 5 and 6 respectively. Results of statistical testing in source space (Bayes factors) are displayed in Table 1.

**Figure 3.**
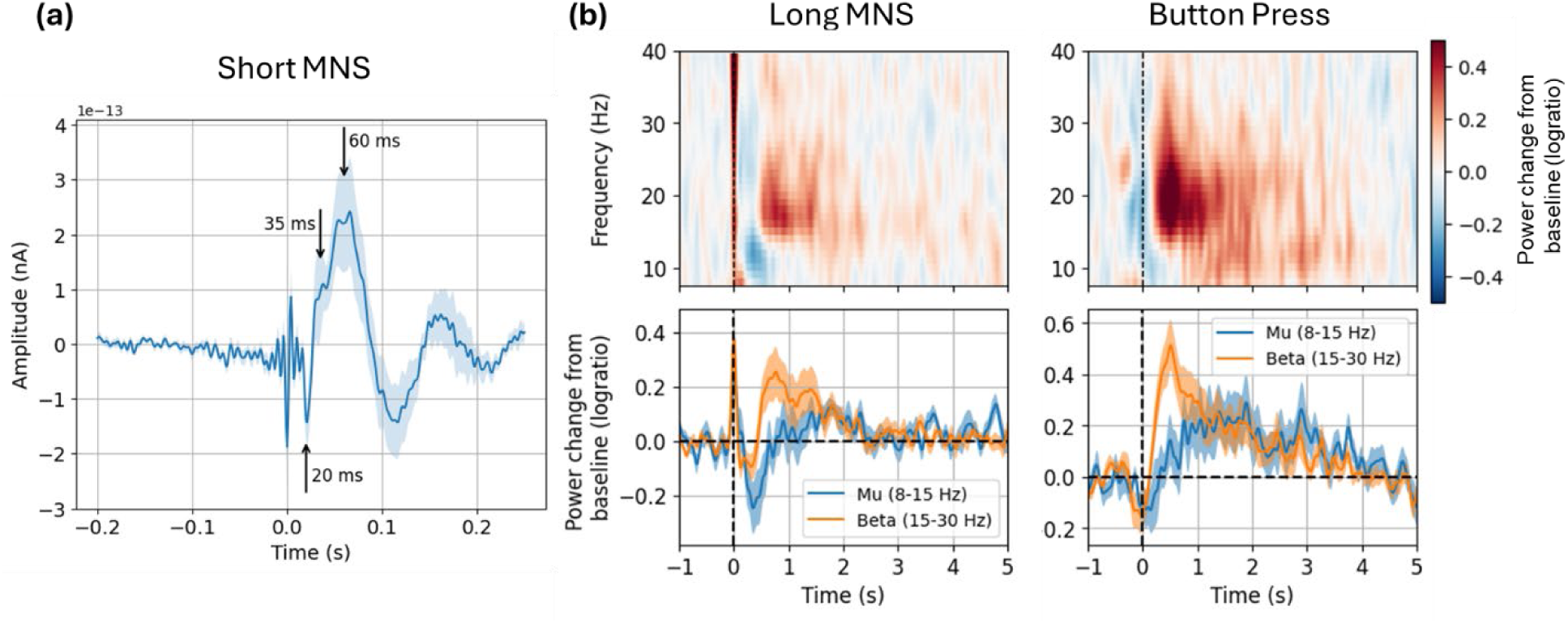
Grand-average results at one sensor (C3). (**a**) The evoked field elicited by the short MNS paradigm; (**b**) Mu and beta modulation elicited by the long MNS paradigm (left) and the button pressing task (right).

**Figure 4.**
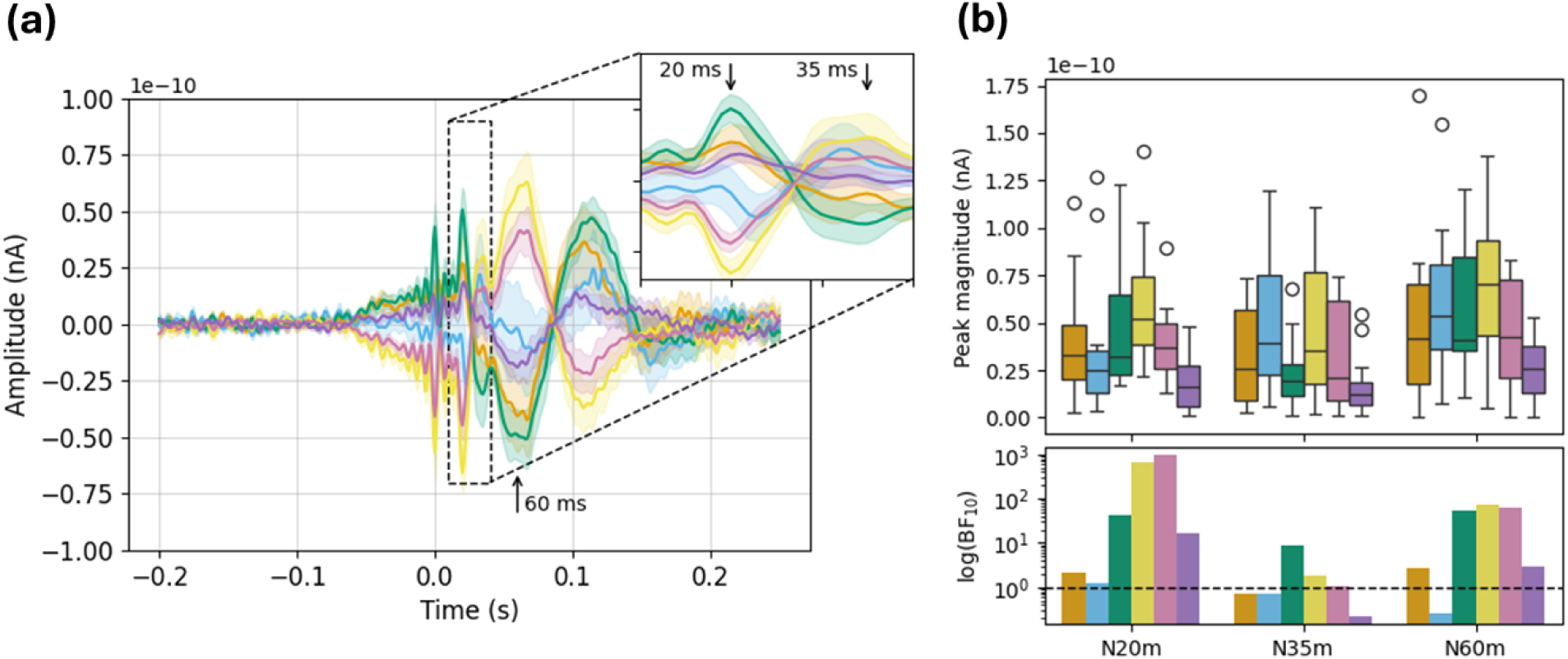
Source-localised SEF results. **(a)** Timecourses show the grand-average evoked field in each ROI; ribbons are +/- 1 SE. **(b)** Top: Boxplots show magnitudes of the N20m, P35m, and P60m peaks (i.e., absolute amplitude difference between peak and baseline windows). Bottom: Bars show Bayes factors for each peak (note: logarithmic y-scale).

**Figure 5.**
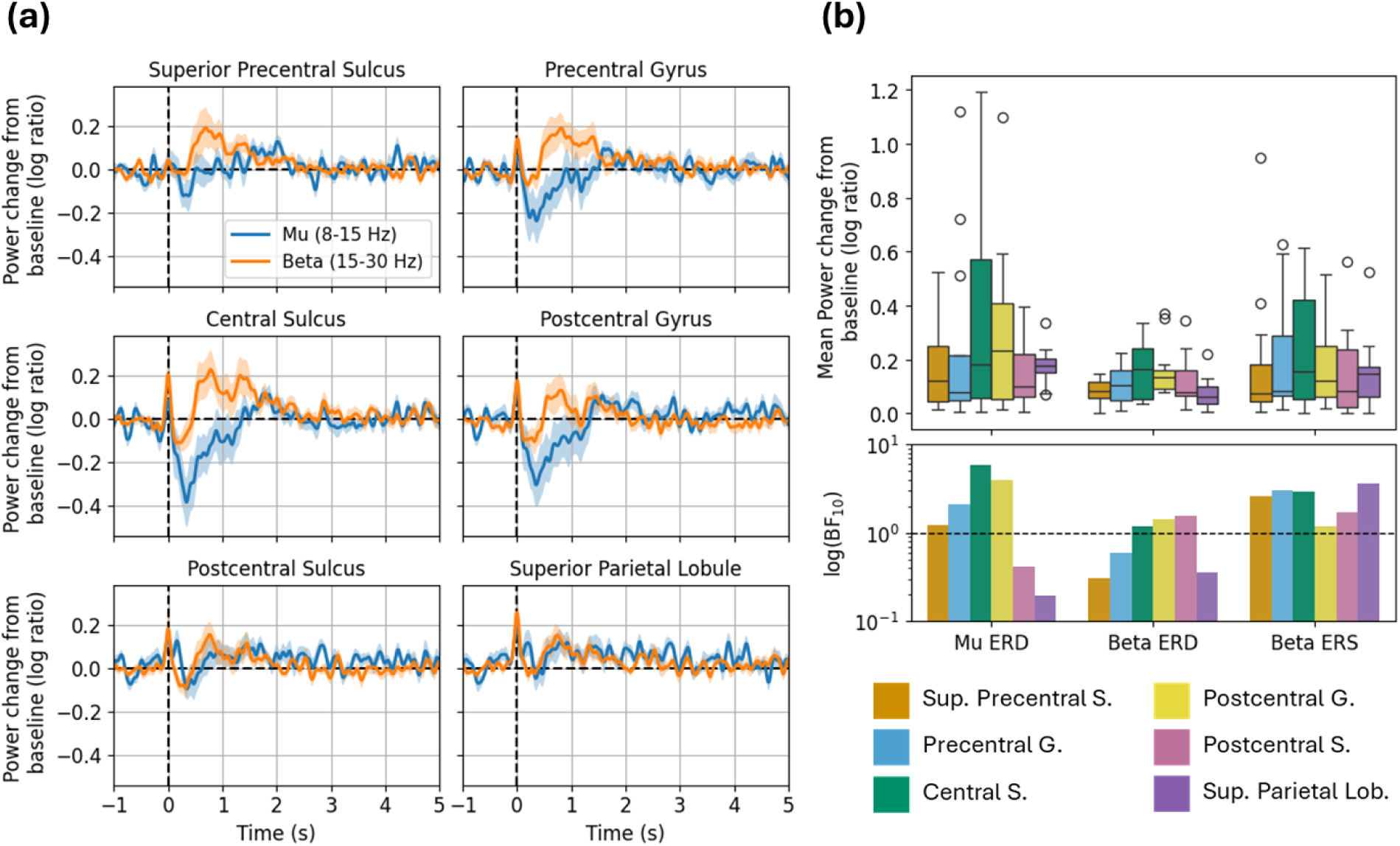
Source-localized mu and beta modulation in long MNS task. **(a)** Grand-average timecourses show power change from baseline in each frequency band; ribbons are ±1 SE. **(b)** Top: Boxplots show mean power change from baseline within each window of interest (mu ERD, beta ERD, and beta ERS). Bottom: Bars show Bayes factors for each window of interest (logarithmic y-scale). Abbreviations: *Sup.* Superior; *S.* sulcus; *G*. gyrus.; *Lob.* Lobule.

**Figure 6.**
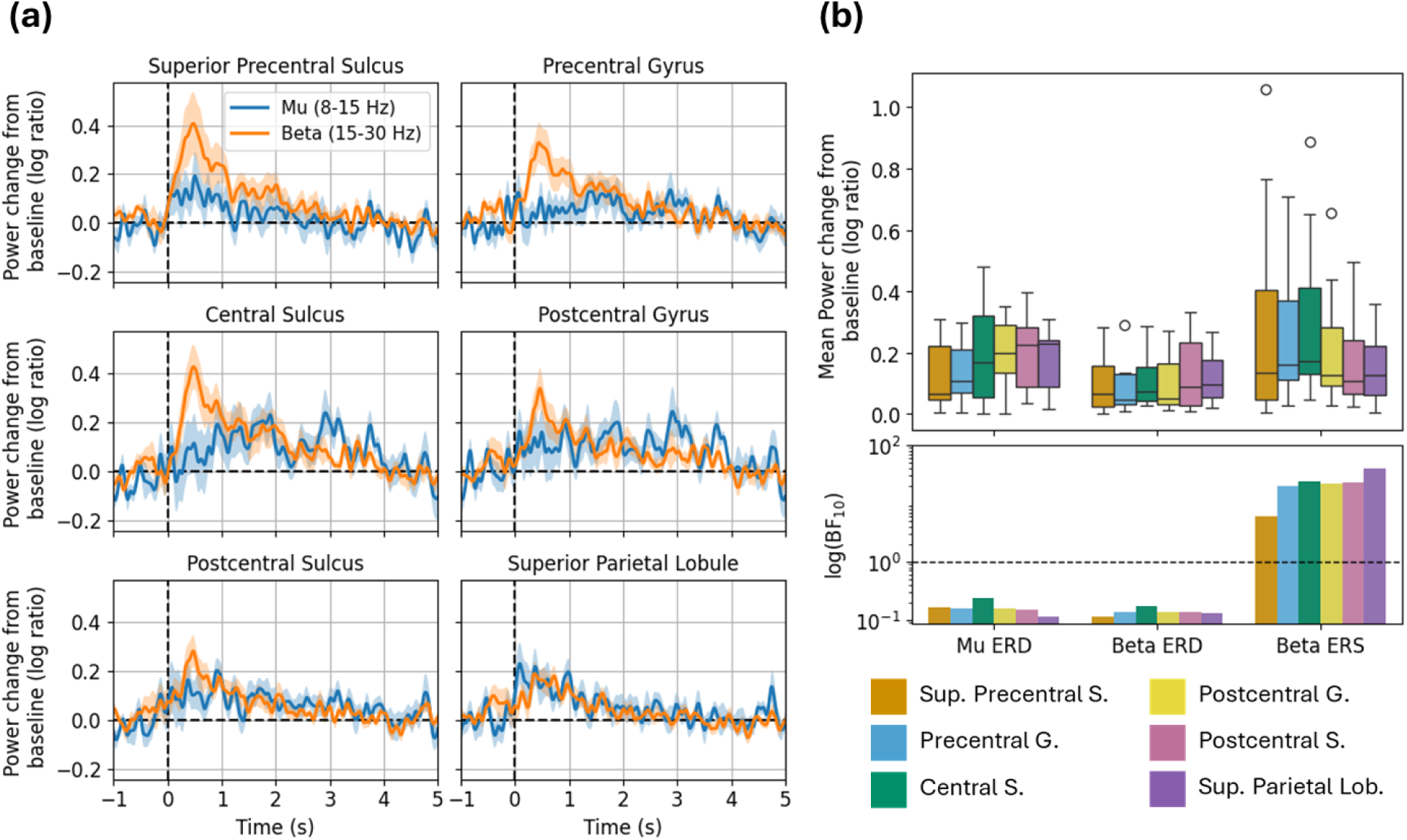
Source-localized mu and beta modulation in the button press task. **(a)** Grand-average timecourses show power change from baseline in each frequency band; ribbons are ±1 SE. **(b)** Top: Boxplots show mean power change within each window of interest (mu ERD, beta ERD, and beta ERS). Bottom: Bars show Bayes factors for each window of interest (logarithmic y-scale). Abbreviations: *Sup.* Superior; *S.* sulcus; *G*. gyrus.; *Lob.* Lobule.

**Table 1.**
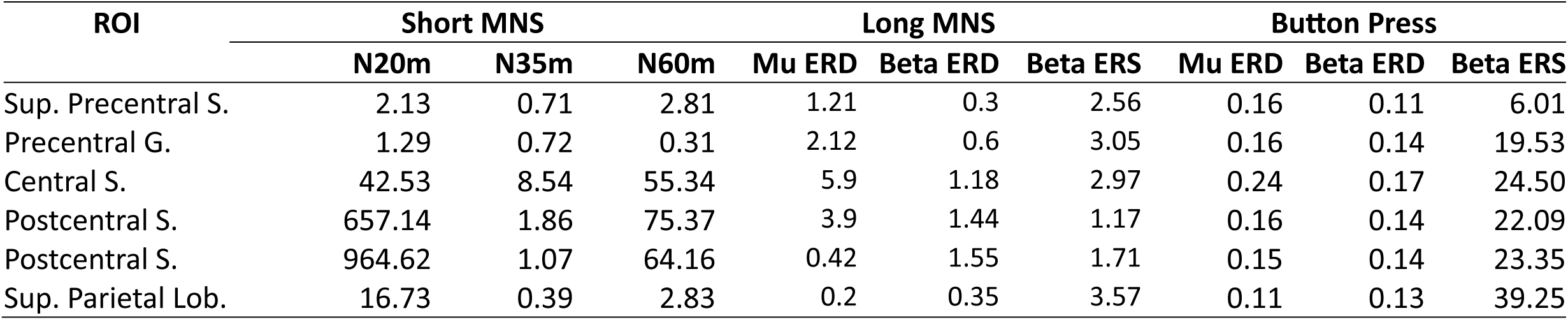
Values are Bayes factors (BF_10_) for source-localized SEF peaks in the short MNS data, and mu/beta rebound peaks in the long MNS and button press data. Abbreviations: *Sup.* Superior; *S.* sulcus; *G*. gyrus.; *Lob.* Lobule.

| ROI | Short MNS |  |  | Long MNS |  |  | Button Press |  |  |
| --- | --- | --- | --- | --- | --- | --- | --- | --- | --- |
|  | N20m | N35m | N60m | Mu ERD | Beta ERD | Beta ERS | Mu ERD | Beta ERD | Beta ERS |
| Sup. Precentral S. | 2.13 | 0.71 | 2.81 | 1.21 | 0.3 | 2.56 | 0.16 | 0.11 | 6.01 |
| Precentral G. | 1.29 | 0.72 | 0.31 | 2.12 | 0.6 | 3.05 | 0.16 | 0.14 | 19.53 |
| Central S. | 42.53 | 8.54 | 55.34 | 5.9 | 1.18 | 2.97 | 0.24 | 0.17 | 24.50 |
| Postcentral S. | 657.14 | 1.86 | 75.37 | 3.9 | 1.44 | 1.17 | 0.16 | 0.14 | 22.09 |
| Postcentral S. | 964.62 | 1.07 | 64.16 | 0.42 | 1.55 | 1.71 | 0.15 | 0.14 | 23.35 |
| Sup. Parietal Lob. | 16.73 | 0.39 | 2.83 | 0.2 | 0.35 | 3.57 | 0.11 | 0.13 | 39.25 |

### Sensor-Level Results

#### SEFs

The grand-average evoked field at one representative sensor (C3), shown in Figure 3a, exhibited the expected SEF complex. It comprised a clear negative peak at 20 ms (corresponding to the canonical N20m) followed by a broad positive deflection which briefly plateaued around 35 ms (P35m) and peaked at 60 ms (P60m).

#### Mu and beta modulation

Figure 3b shows grand-average time-frequency responses (TFRs), as well mu and beta timecourses, at the same sensor. The long MNS data shows the expected suppression (ERD) in the mu and beta frequency ranges immediately following stimulation (with greater in magnitude in the mu range), followed by ERS in the beta range after around 0.5 s. A brief period of marginal amplitude mu and beta ERD was evident in the button-press data, and there was a clear ERS effect in the beta range after 0.5 s.

### Source-Level Results

#### SEFs

The grand-average SEF for each ROI in the short MNS paradigm is shown in Figure 4a. Here we saw clear peaks at 20 and 60 ms (and 35 ms to a lesser extent), particularly in the central and postcentral sulci, and the postcentral gyrus. Magnitudes for each peak were calculated as the absolute amplitude difference between each 3 ms time window and the baseline window (see Methods). With respect to the N20m and P60m: these peaks were maximal in the postcentral gyrus and decreased linearly in the anterior and posterior directions (Figure 4b). Results of our statistical analyses showed a similar trend: we saw the strongest evidence for the N20m and P60m (BF_10_ values around 600-1000 and 65-75 respectively) in the postcentral gyrus and sulcus, with strength of evidence decreasing in the anterior and posterior directions. These results are consistent with a neural generator in primary somatosensory cortex. Results for the P35m were more equivocal—statistical evidence for this peak was weak (compared with the N20m and P60m), and we did not see a clear spatial trend in peak magnitudes. We therefore consider the N20m and P60m to be robustly detected; the P35m less so.

#### Mu and beta modulation

In the long MNS data (Figure 5), the expected mu ERD and beta ERS effects were evident in multiple ROIs. Mu ERD was most evident in the central sulcus and postcentral gyrus; mean power change and strength of evidence was maximal in this ROI and decreased in the anterior and posterior directions. We saw a similar trend with respect to beta ERD, though in this case the statistical evidence was relatively weak. Meanwhile, beta ERS appeared more anteriorly to the ERD effects: broadly, we saw higher power change and strength of evidence in the superior precentral sulcus, precentral gyrus, and central sulcus compared to other ROIs.

In the button press data (Figure 6), we did not see the expected mu/beta ERD effects in any ROIs (BF_10_ values were consistently < 1.0, indicating support for H_0_ over H_1_); however there was strong evidence for beta ERS. In terms of source strength, the beta ERS effects followed the same spatial pattern to that of the long MNS data (i.e., higher power change in anterior ROIs); however the statistical results indicated stronger evidence in central and posterior ROIs. In contrast to the long MNS results, this suggests are more central/posterior neural generator for beta ERS.

We note that, in both the long MNS and button press data, we observed the strongest evidence for beta ERS in the superior parietal lobule, despite the actual magnitude of this effect being relatively small in both cases (Figures 5b, 6b). This finding is unexpected, and seems at odds with broader numerical and statistical trends described above. We return to this issue in the Discussion.

### Signal-To-Noise Ratio (SNR)

We computed sensor- and source-level SNR for the N20m, for each participant. We focused on the N20m due to the ubiquitous and robust nature of this response in MEG. At the sensor level, mean and median SNR across subjects was 8.66 and 7.41 respectively (range: 3.05-21.45), and we identified C3 as the most common sensor location at which SNR was maximal (C3 yielded the best SNR in 25% subjects). At the source level mean and median SNR were 8.67 and 8.66 (range: 1.22-25.87). The central sulcus was the most common ROI at which SNR was maximal (33.33% of subjects).

## Discussion

In this study we sought to validate the use of a mobile, low-cost OPM-MEG system— comprising a cylindrical mu-metal shield and partial head coverage sensor array—for recording neuromagnetic responses in humans. To this end, we investigated evoked and oscillatory sensorimotor responses (which are robustly detected by other OPM-MEG systems; see Introduction) following median nerve stimulation (MNS) and voluntary movement. These included the N20m, P35m, and P60m components of the somatosensory evoked field (SEF) complex, and task-related modulation of mu and beta rhythms (beta and mu ERD; beta ERS). We investigated these responses in three datasets, acquired during two MNS paradigms and a cued button-pressing task. Furthermore, we assessed the spatial distribution of these effects on the cortical surface, as revealed by distributed source modelling. To facilitate direct comparison to previous MEG work, we also computed sensor- and source-level SNR around the N20m peak.

Broadly speaking, our sensor- and source-level results from the MNS paradigms were consistent with our predictions. With respect to the SEF complex, we identified clear peaks at 20 and 60 ms following stimulation, corresponding to the canonical N20m and P60m. At the source level these peaks were maximal (and accompanied by the strongest statistical support) in the postcentral gyrus and adjacent ROIs. This finding is consistent with previous localizations of SEF peaks to primary somatosensory cortex (Lin et al., 2005; Ou et al., 2009; Wikström et al., 1996). Our ability to isolate the P35m is likely limited by the relatively low sample rate and bandwidth of our system; most investigations of early SEF peaks acquire data at 1000’s Hz with low-pass filtering at half the sample rate. We also observed the expected mu (and, to a lesser extent, beta) ERD immediately following stimulation, and a subsequent beta rebound (ERS) effect. At the source level, ERD and ERS were most evident in central-posterior and central-anterior ROIs respectively, again consistent with previous work (Bardouille et al., 2019; Jurkiewicz et al., 2006). These findings provide encouraging proof-of-principle support for our system. Not only could we detect these canonical responses; they also conformed to expected spatial distributions along the cortical surface. We note that previous work with a similar cylindrical-shield system has reported SEF and oscillatory sensorimotor responses at the sensor level (Borna et al., 2017), and successful localisation of SEFs with dipole fitting (Borna et al., 2020; Boto et al., 2017). Our MNS results provide convergent support for these findings, and further demonstrate the viability of distributed source modelling approaches in such systems.

Results from the button press task did not conform to our expectations. While we did detect beta ERS at both the sensor and source level, the expected ERD effects were absent. Moreover, at the cortical level, evidence for beta ERS following button pressing was weak in anterior ROIs and relatively strong in posterior ROIs—contrary to our MNS results. The discrepancy is unlikely due to participant sampling or statistical power (all data were collected from the same participants, within the same scanning session, and the two tasks comprised the same number of trials). It seems, therefore, that some component of the button press task interfered with our ability to accurately characterise mu/beta modulations in this system. One possibility is that button presses induced movement of participants’ heads relative to the (fixed) sensor array, which would disrupt data quality. However, this seems unlikely since button-press tasks are commonplace in fixed-array cryogenic systems (e.g., Barratt et al., 2017; Embury et al., 2019; Schmiedt-Fehr et al., 2016; Taylor et al., 2017); moreover, we did not observe such movement artefacts in the raw data. An alternative explanation is some interplay between the active nature of the task and the sensitivity of OPMs to vibration-induced noise that overlays the beta band. For example, participant movement (such as a button press) induces small vibrations in the scanner bed (to which the OPM helmet was affixed) leading to vibration-induced field changes. One must also consider that effective neuroimaging in active tasks (e.g., button pressing) relies on appropriate participant engagement more so than passive tasks (e.g. MNS). Given the small sample size, it is possible that errant participant behaviour may be responsible for this surprising discrepancy in findings. Future work might explore modified apparatus (e.g., cushioning under the response box), concurrent electromyography recording, or alternative paradigms (e.g., cued finger flexion, as in Labyt et al., 2003; Pfurtscheller & Neuper, 1997; Salmelin & Hari, 1994; Wilson et al., 2010) to explore voluntary movement with this system.

We also computed signal-to-noise ratio (SNR) around the N20m peak for direct quantitative comparison to other MEG systems. Previous work has reported N20m SNR in the range of around 12-48 in SQUID-MEG systems (Antonakakis et al., 2020; Borna et al., 2020; Boto et al., 2017; Kimura & Hashimoto, 2001). With respect to OPM-MEG, previous work with partial-coverage arrays has reported SNRs around 42 at the source level (Boto et al., 2017) and 14-25 at the sensor level (Borna et al., 2017) (we are not aware of any studies reporting N20m SNR in whole-head OPM). By comparison, our system yielded relatively low SNR: around 7-8 (median) at both the sensor and source levels, with a maximum of 25.87 in one participant. A few factors likely contributed to this apparent reduction in SNR. First, our use of a small and passive shield likely provided less attenuation of environmental noise compared with conventional MSRs (used in Antonakakis et al., 2020; Boto et al., 2017; Kimura & Hashimoto, 2001), which typically benefit from both passive shielding and active field nulling. Indeed, Borna et al., 2020 also employed active nulling in their cylindrical shield. Moreover, as noted earlier, our system was limited to a sample rate of 1000 Hz. This is considerably lower than the sample rate of other investigations of the N20m (often >= 10 KHz, e.g., Borna et al., 2020; Boto et al., 2017). Given the temporal proximity of the stimulus artifact and N20m, the lower sample rate likely further impacted SNR. Finally our limited head coverage, combined with our use of single-axis sensors, limited our ability to attenuate noise post-hoc. In the context of source localization, noise suppression tends to increase with channel count (Vrba et al., 2004). Similarly, homogenous field correction (HFC; an important noise reduction step applied during preprocessing) is more effective when applied to data with a plurality of sensor orientations (Tierney et al., 2021). In light of these considerations, it is not too surprising that we observed relatively low SNR compared with other systems. Future improvements—in particular, incorporating multi-axis sensors and/or hardware capable of higher sampling rates—will likely yield SNR closer to that of other systems.

Another consideration is that we did not establish, a-priori, the spatial extent of our effective sensitivity on the cortical surface, given our sensor count and layout. As such, it is not clear whether all our ROIs lay within the area from which we could reliably detect cortical responses. Our sensor array appears to have provided good coverage of primary sensorimotor cortices, given our results in those ROIs. However, it is possible that we had poorer sensitivity to the more peripheral ROIs. This may help to contextualize the strange finding that the superior parietal lobule (an area not typically associated with sensorimotor processing) exhibited the strongest evidence for beta ERS, despite very low source estimated response amplitude, in two datasets. This area also showed consistently (and suspiciously) low variance in all three datasets, evidenced by markedly narrower error bars and boxplots, compared with other ROIs (Figures 4b-6b). We do not have a clear explanation for this low variance, but it may reflect a spatial coverage problem. Future work with partial-coverage systems—particularly those with low sensor counts—should aim to establish optimal sensor layouts for targeting specific areas. One solution is to simulate dipoles in target locations and determine sensitivity at a given set of sensor locations (e.g., illustrated in Hill et al., 2022 Figure 1b). In a related vein, Hill et al. (2024) recently demonstrated that source-level SNR can be improved through optimal sensor arrangement. Future work might characterise the spatial extent of sensitivity at the cortical surface (perhaps at some SNR threshold) based on varying sensor layouts with a fixed number of sensors at known orientations. This would allow researchers to determine the optimal layout for targeting a specific set of areas, given a limited number of sensors.

## Conclusions

Our study provides preliminary support for a mobile, low-cost OPM-MEG system for human recording, and adds to the growing body of work seeking to develop more accessible and flexible OPM solutions (Bardouille et al., 2024; Borna et al., 2017, 2020; Hill et al., 2024; Holmes et al., 2022, 2023; Schofield et al., 2024). Our system was able to detect and localize canonical neuromagnetic responses in passive paradigms such as MNS (and, potentially, resting-state scans), though further work is needed to validate its use in the context of voluntary movement.

Ongoing developments in mobile OPM scanners (Bardouille et al., 2024; Borna et al., 2017, 2020) show promise for a viable alternative to modern state-of-the-art systems housed in MSRs. At present, such systems offer superior SNR, likely owing to high channel counts and active field control. However, we have argued that improvements to our mobile scanner (such as incorporating multi-axis sensors, field nulling, and increasing sampling rate), as well as determining optimal sensor layouts a-priori, could potentially bring our SNR to comparable levels. We hope that ongoing work in this area will open the door to a range of applications. For example, OPM has shown promise for detecting and localizing focal epilepsy (Vivekananda et al., 2020) and traumatic brain injury (Dunkley, 2022); validation of mobile and inexpensive scanners could potentially improve clinical accessibility, particularly in the contexts of small community and mobile healthcare delivery. Efforts to expand OPM to these and other applications will naturally lead to further validation and refinement of this technology.

## List of Abbreviations

ERD: Event-related desynchronization
ERS: Event-related synchronization
MEG: Magnetoencephalography
MNS: Median nerve stimulation
MSR: Magnetically shielded room
OPM: Optically-pumped magnetometer
SEF: Somatosensory evoked field
SNR: Signal-to-noise ratio
SQUID: Superconducting quantum interference device

## Acknowledgements

We would like to thank Clara Knox and Carson Leslie for assisting with data collection in this study.

## Code Availability Statement

Code for all analyses presented in this manuscript is publicly available on GitHub: https://github.com/lyambailey/OPM_MNSBP.

## Ethics Statement

All procedures were approved by the Dalhousie University Research Ethics Board (REB #2021-5586).

## CRediT Author Statement

**Lyam M. Bailey:** Conceptualization, Data curation, Writing - original draft, Formal analysis, Investigation, Methodology, Software, Validation, Visualization. **Timothy Bardouille:** Conceptualization, Data curation, Funding acquisition, Project administration, Methodology, Resources, Software, Validation, Visualization, Supervision, Writing - review & editing, Formal analysis.

## Footnotes

1 Left handed individuals were not deliberately excluded from this study, as per Bailey et al. (2019).

2 This was achieved by supplying the *pca_flip* option to the *mode* argument for the *extract_label_timecourses()* MNE function, see: https://mne.tools/stable/generated/mne.extract_label_time_course.html

3 We did not have a-priori predictions about the polarity of each peak in source space (since polarity depends on orientation of the underlying source, and therefore will differ between ROIs). However, the polarity of the P35m and P60m should be opposite to that of the N20m in all ROIs. Therefore, we determined the direction (left- or right-sided) of our one-sampled t-tests based on the polarity of the N20m in the grand-average evoked field, independently for each ROI. For example, if the N20m was negative in a given ROI then we used a left-sided test to evaluate the N20m, and right-sided tests to evaluate the P35m and P60m. Using one-sided tests here ensured equivalent statistical stringency between the SEF analysis and the oscillatory analyses.

4 All t-tests used half-Cauchy JZS priors. JZS priors are default in the BayesFactor package (Morey et al., 2022); the only change we made to default parameters was using a half-Cauchy (rather than full) to enable one-tailed tests.

